# Recombinant *Ehrlichia* TRP75 Rescues Molting Failure of Ticks Infected with TRP75 Mutant *Ehrlichia*

**DOI:** 10.64898/2026.10.26.761916

**Authors:** Jeffrey Kawahara, Tsian Zhang, Lidan Wang, Stephen L. Denton, Mingqun Lin, Yasuko Rikihisa

## Abstract

*Ehrlichia* species are obligatory intracellular bacteria that exist in nature through a cycle of infection and transmission between ticks and their mammalian hosts, potentially causing a severe febrile illness called ehrlichiosis in humans and animals. Roles of *Ehrlichia* molecules in tick-mammal *Ehrlichia* transmission are mostly unknown. Tandem repeat protein (TRP) 75 is a lipoprotein conserved among *Ehrlichia* species. We identified and cloned a Himar1 transposon insertional mutant of *Ehrlichia japonica* that lacks C-terminal intrinsically disordered region of TRP75 (TRP75ΔC). This mutant was competent in infecting and replicating in both mammalian and tick cells in culture, and causing fatal illness similar to wild-type (WT) *E. japonica*. Upon feeding on TRP75ΔC and WT *Ehrlichia*-infected mice, both larval- and nymphal-stage of *Ixodes scapularis* ticks ingested similar amounts of blood meal as those fed on uninfected mice, and acquired similar levels of infection. However, most of TRP75ΔC *Ehrlichia*-infected ticks failed to complete molting, in contrast to successful molting of WT *Ehrlichia*-infected and uninfected ticks by 10 weeks post-feeding. In both WT *Ehrlichia*-infected and uninfected ticks, ∼1,000-fold exponential increase in ecdysone receptor (EcR) mRNA occurred at 15 days post-feeding, whereas EcR mRNA was not upregulated in TRP75ΔC *Ehrlichia*-infected ticks. Upon microinjection of recombinant full-length TRP75 protein into engorged ticks infected with TRP75ΔC *Ehrlichia,* molting efficiency was ∼90% restored, and there was significantly increased EcR mRNA compared with solvent microinjection control. This suggests that the endoparasitic bacterial molecule TRP75 manipulates the molting success of infected ticks and EcR mRNA levels for the survival of bacteria-hosted ticks and transstadial transmission.

**Importance:** Tick-borne diseases represent an increasing threat throughout the world. Among these diseases are those caused by obligatory intracellular bacteria of the order Rickettsiales including members of *Ehrlichia* species. Effective prevention of tick-borne diseases requires an understanding of how disease-causing organisms thrive in ticks and are transmitted to mammals. Here, we used our mouse model of *Ehrlichia japonica* tick transmission to investigate the role of a single *Ehrlichia* factor, namely TRP75, using an isogenic Himar1 transposon mutant of TRP75 to infect mammalian and tick hosts and study transmission. Our results demonstrate that full-length TRP75 is necessary to permit molting of *Ehrlichia-*infected ticks via ecdysone receptor upregulation. The results provide new insights into the importance of bacterial factors in altering the molting and survival of infected ticks by allowing ecdysone signaling, thereby promoting pathogen survival and transmission.

## INTRODUCTION

The prevalence of tick-borne diseases, including those caused by obligatory intracellular bacteria such as *Ehrlichia spp*., is increasing worldwide (1–3). The etiological agent of human monocytic ehrlichiosis (HME), *Ehrlichia chaffeensis*, is a classic example of this class of emerging pathogens. HME presentation varies from asymptomatic infection to acute severe morbidity and death (2-5% mortality) (4–7). Although doxycycline is generally effective in treating HME (5), no alternatives exist should microbial resistance develop. No effective vaccine is currently available for any ehrlichiosis of humans or animals.

Ticks acquire ehrlichiae by biting infected vertebrates. Then, ehrlichiae are transstadially transmitted through molting to the next developmental stage of ticks (larva → nymph → adult) prior to transmission to a next vertebrate host, including humans (8). *E. japonica* (formerly, *Ehrlichia* sp. HF, *Ixodes ovatus Ehrlichia* [IOE] agent) (9), was originally isolated from *Ixodes ovatus* ticks, but also has been found in rodents, shrews, and dogs in Japan (10–13). *E. japonica* has a relatively broad geographic distribution and several tick vectors, and indeed it has been found in *Ixodes ricinus* and *I. apronophorus* ticks in several European countries (14–16). *E. chaffeensis* is most closely related to *E. japonica* (11, 17). Laboratory mice infected with *E. japonica* experience progressive infection that may lead to the subclinical-to-fatal disease in a dose-dependent manner (LD_50_ = 100 bacteria) (18), and the immunopathogenesis of which has been studied extensively (11, 19–25). As there is no small animal model of disease or tick transmission of *E. chaffeensis*, *E. japonica* serves as a valuable component of a small animal disease model of human ehrlichiosis and tick transmission.

Although several unique ehrlichial molecules in addition to house-keeping genes are required for infection of mammalian or tick cells in culture (26–28), or mice (18, 28, 29), the ehrlichial molecules that drive the tick-mammal transmission are largely unknown. Following successful cultivation of *E. japonica* in DH82 canine histiocytic leukemia cell line and the subsequent achievement of whole-genome sequencing (17), we constructed a Himar1 transposon mutant library comprising at least 158 distinct *E. japonica* mutants (18). We recently used one of the cloned isogenic Himar1 transposon knockout mutants to demonstrate that tandem repeat protein (TRP)120 is required for tick acquisition of *Ehrlichia* species from infected mice (30). We further developed an additional 249 distinct mutants (31). Among these new Himar1 transposon mutants is *E. japonica* EHF_0506 (*TRP75*)::Himar1, encoding an insertional mutant of TRP75. Based on BLAST analysis, the expected E-value between *E. chaffeensis* and *E. japonica* TRP75 is 6e^-149^ (identity: 62.3%; similarity: 75%), demonstrating that two homologs are highly similar in amino-acid sequence (32). *E. chaffeensis* TRP75 is a lipoprotein (33) and is reported to be a possible effector of the Type I secretion system, and predicted to interact with 90 different human proteins, suggesting that TRP75 alters human cell pathophysiology (34, 35). However, hitherto, nothing is known about the functions of TRP75 during *Ehrlichia* infection of tick cells in culture, tick and mammalian host infection, and tick transmission *in vivo*.

To address these issues, we first cloned and characterized the *E. japonica TRP75*::Himar1 mutant and investigated its ability to infect cultures of mammalian cells and tick cells in culture, and its infection and disease in mice. Further, we used a mouse model of *Ixodes* tick *Ehrlichia* transmission (30) to examine the roles of TRP75 in tick infection with *Ehrlichia* for its transmission between mammals and ticks. Our results indicate that C-terminal intrinsically disordered domain (IDR) of TRP75 is the first example of an *Ehrlichia* factor that is required for infected tick molting, thus required for survival of both *Ehrlichia* and its hosting tick. Consequently, TRP75 IDR is required for transmission from infected ticks to mammals, despite TRP75 IDR is dispensable for mammalian or tick cell infection and *in vivo* mouse pathogenesis.

## RESULTS

### Cloned *TRP75*::Himar1 *E. japonica* has a single Himar1 insertion per genome and produces a C-terminally truncated TRP75 protein in infected cells

*E. japonica* HF TRP75 comprises 684 amino acids (aa) encoded by EHF_0506 (2,055 bp) and contains 17 tandem repeats of 22-aa each (Fig. 1A). In the *TRP75*::Himar1 mutant, the 1.8 kb Himar1 sequence containing mCherry and the selection marker for spectinomycin/streptomycin resistance genes were mapped at genomic locus 556,130, which is downstream of the TRP repeats (near the 3’-end) of *TRP75* gene (Fig. 1A). To characterize this mutant, *TRP75*::Himar1 mutant was cloned from the original culture designated H130-A1 containing *TRP75*::Himar1 mutant by repeated limited dilution and passages in DH82 cell culture, followed by multiple passages of the cell culture-cloned mutant in mice as previously described (18, 30). Cloned *TRP75*::Himar1 was verified by insertion site–specific “flanking PCR” (18) for *TRP75* gene, showing a single larger PCR product containing the Himar1 insert (Fig. 1B). Himar1 insertions occur mostly once per genome, and rarely twice (36, 37). To verify that there were no additional Himar1 insertions in the genome of cloned *TRP75*::Himar1, quantitative PCR (qPCR) was performed and the results showed an approximate ratio of 1:1 for the copy numbers of the mCherry gene vs. the single-copy *Ehrlichia* 16S rRNA gene (Fig. 1C), indicating a single Himar1 transposon insertion per *Ehrlichia* genome. *TRP75*::Himar1 *E. japonica* replicated well in DH82 (canine macrophage cell line), RF/6A (monkey retinal endothelial cell line), and ISE6 (*Ixodes scapularis* tick embryonic cell line), and several large bacteria-containing morulae (intracellular microcolonies of bacteria) are present in the cytoplasm similar to observation with WT *E. japonica* (Fig. 1D).

**FIG 1.**
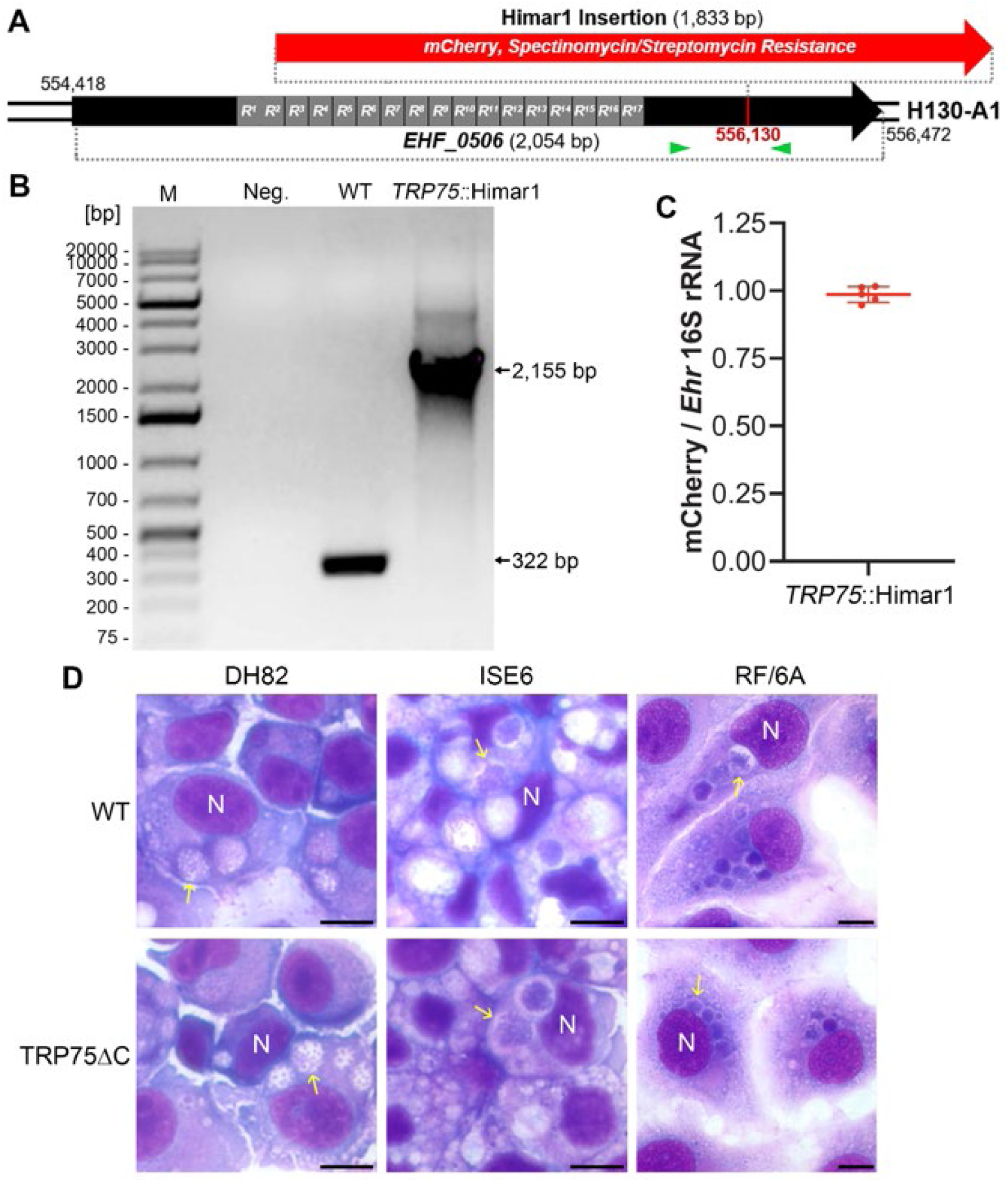
Cloned *TRP75*::Himar1 mutant of *E. japonica.* (**A**) Himar1 transposon insertion site of mCherry and spectinomycin/streptomycin resistance genes in *E. japonica TRP75 (EHF_0506)*. The insertion site in the clonal H130-A1 isolate was mapped at the genomic locus of 556,130, downstream of 17 tandem repeats of 22 aa (gray boxes, R1 – R17). (**B**) Himar1 transposon insertion site–specific flanking PCR to verify mutant clonality. Primer locations are indicated with green arrows in panel A. M, DNA ladder in base pair (bp). PCR product size for WT was 322 bp, and *TRP75*::Himar1 was 2,155 bp with the Himar1 insertion sequence (1,833 bp). (**C)** The ratio of copy numbers of mCherry vs. *Ehrlichia* (*Ehr*) 16S rRNA. DNA samples were purified from five independent passages of DH82 cells infected with the same clone of *TRP75*::Himar1 *E. japonica*, and qPCR was performed with primers for mCherry and *Ehrlichia* 16S rRNA gene. (**D**) Mammalian cells (DH82 or RF/6A) or tick cells (ISE6) infected with WT or *TRP75*::Himar1 *Ehrlichia*. HEMA 3 staining. Examples of bacteria-containing morulae were indicated by yellow arrows. N, nucleus. Bar, 10 µm.

The Himar1 insertion in TRP75 created a premature stop codon downstream of the transposon inverted terminal repeat (ITR) sequence, truncating TRP75 C-terminal 113 aa (Fig. 2A). Hence, we named this *E. japonica* mutant as TRP75ΔC. By Western blotting using dog antiserum developed against recombinant full-length *E. chaffeensis* TRP75, both *E. japonica* and *E. chaffeensis* native TRP75 were detected at ∼90-100 kDa (Fig. 2B), a range that was larger than the predicted molecular mass (*E. japonica* 78.2 kDa, *E. chaffeensis* 66.2 kDa), whereas *E. japonica* TRP75ΔC was ∼ 80 kDa (predicted molecular mass, 66.0 kDa). The larger size in each case was unlikely attributable to post-translational modifications of *Ehrlichia* TRP75, such as glycosylation, SUMOylation, or phosphorylation of Ser/Thr/Tyr, because rTRP75 of *E. chaffeensis* and *E. japonica* purified from *Escherichia coli,* which is incapable of these modifications, similarly appear as a larger size by Western blotting (33) (Fig. 7A). *Ehrlichia* major outer membrane protein P28, which is encoded by an *omp-1/p28* multigene family (38), was expressed similarly between *E. japonica* WT and TRP75ΔC mutant (Fig. 2B). *E. japonica* TRP75 is predicted to possess a lipobox motif, regions typically important for lipoprotein maturation and anchoring in membranes, as does *E. chaffeensis* TRP75 (33) (Fig. 2C). Based on alpha-fold prediction, the N-terminal lipobox and tandem repeat structure were present in TRP75ΔC, but the majority of C-terminal intrinsically disordered region (IDR) was truncated (Fig. 2C). Alignment of these IDR *E. japonica* TRP75 (downstream of Himar1 insertion, 113 aa) with *E. chaffeensis* TRP75 IDR (C-terminal 116 aa) revealed that they are highly conserved (72% similarity) (Fig. S1). By immunofluorescence labeling, WT *E. japonica* TRP75 protein was found to encase *Ehrlichia* nucleoids (revealed by DAPI stained) within morulae of RF/6A cells and also found on the host cell surface as puncta (yellow arrows), whereas TRP75ΔC protein overlapped diffusely with the *Ehrlichia* nucleoid and was found less abundant on the host cell surface (Fig. 3C).

**FIG 2.**
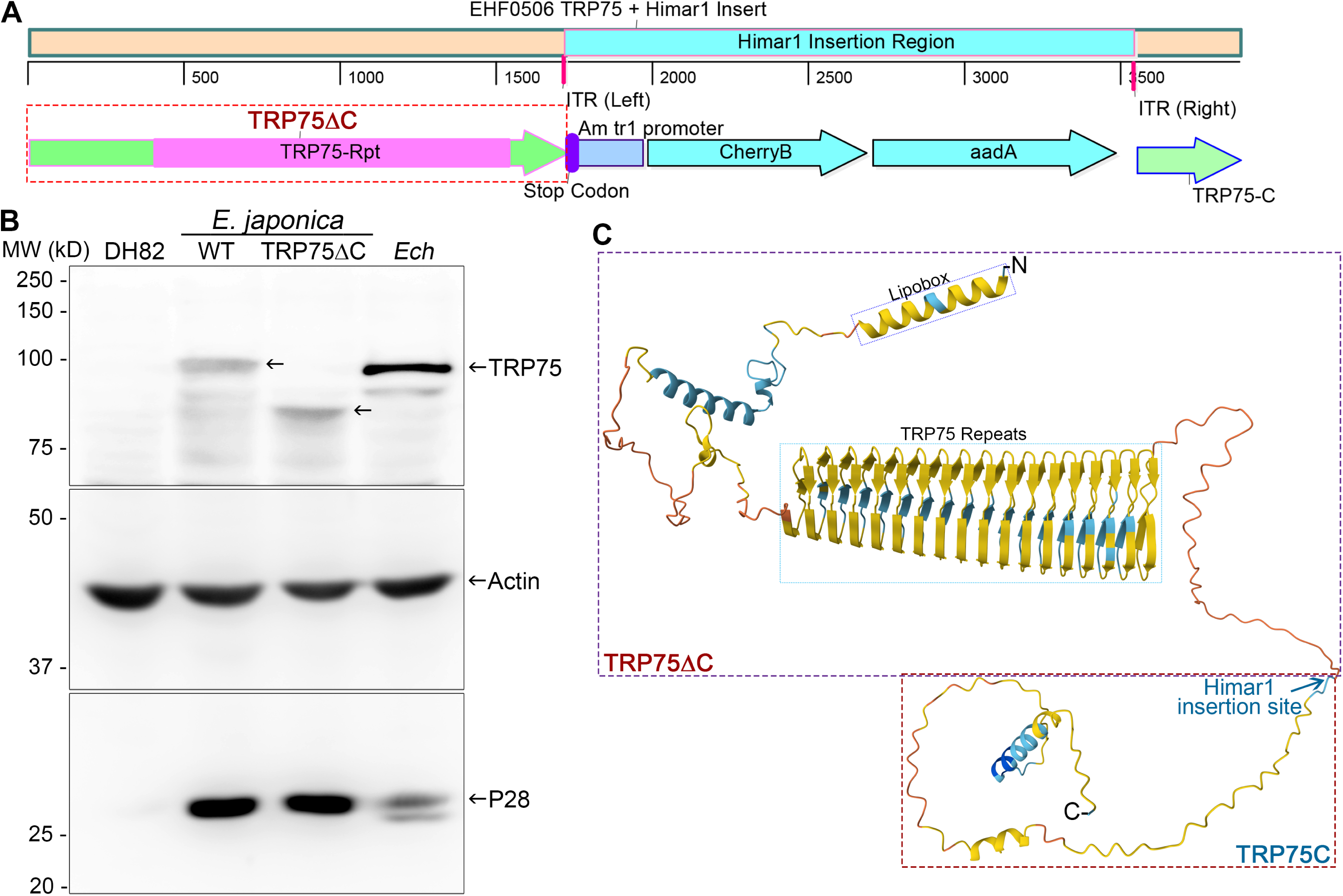
*TRP75*::Himar1 *Ehrlichia* produces a C-terminal truncated TRP75 protein. (**A**) Sequence map of the *TRP75* gene with the Himar1 insertion, which resulted in a premature stop codon (UGA) at the 3’-end of the left inverted terminal repeats (ITR, indicated by the red line) sequence. TRP75-Rpt, tandem repeat domain of TRP75; *aad7*, Spectinomycin/streptomycin resistance cassette; Am tr1 promoter, *A. marginale* tr1 promotor. (**B**) Western blot analysis of WT and TRP75ΔC *E. japonica*-infected, *E. chaffeensis* (*Ech*)-infected, and uninfected DH82 cells labeled with dog anti-*Ech* TRP75, mouse anti-β-actin, and rabbit anti-*Ech* P28, and detected with respective peroxidase-conjugated secondary antibodies. TRP75 protein bands were indicated by arrows in the top blot. (**C**) Predicted three-dimensional structure of TRP75 generated by AlphaFold. Blue arrow indicates Himar1 insertion site, TRP75ΔC and the C-terminus IDR truncated by Himar1 insertion (TRP75C, 113 aa) are indicated in boxes. TRP75C comprises 16.5% of full-length TRP75 (684 aa). Predicted lipobox and TRP75 tandem repeats (17 repeats) within TRP75ΔC are marked by small blue boxes, respectively. Color-coded protein strands indicate confidence metrics shown in pLDDT scores: blue, very high (pLDDT > 90); light blue, confident (90 > pLDDT > 70); yellow, low (70 > pLDDT > 50); orange, low (pLDDT < 50).

**FIG 3.**
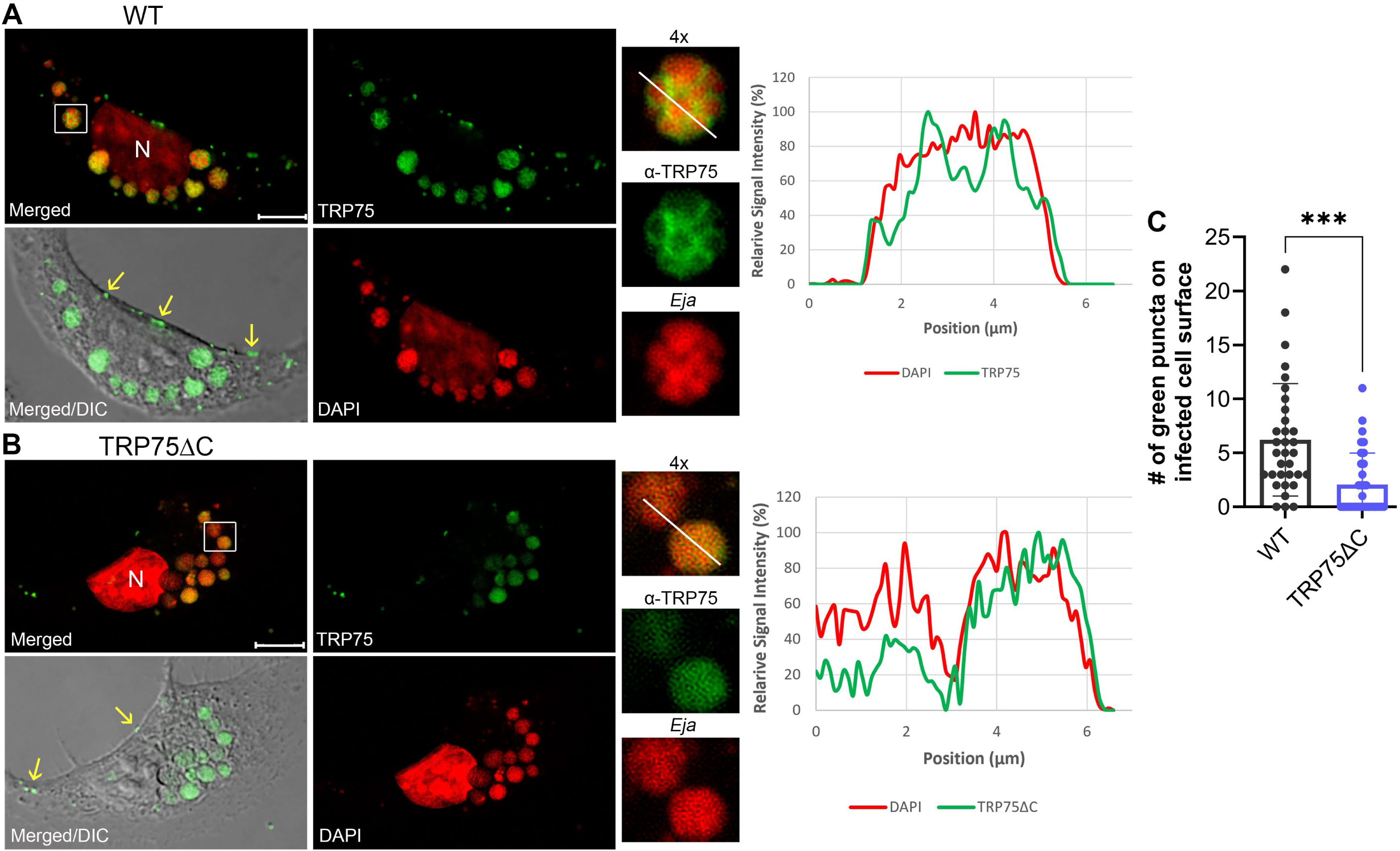
Localization of native TRP75 and TRP75ΔC in infected cells. WT (**A**) or TRP75ΔC (**B**) *E. japonica* (*Eja*)-infected RF/6A cells at 2 dpi were fixed and labeled with dog anti-TRP75 and AF488-conjugated anti–dog IgG. Host cell nuclei (N) and *Eja* DNA were labeled with DAPI (pseudocolored red). DIC, differential interference contrast; merged, fluorescence images merged. Green puncta on the surface of infected cells were indicated by yellow arrows. Each boxed area is enlarged 4× on the right. Scale bars, 10 μm. Relative fluorescence signal intensity plots along the yellow lines in the enlarged merge images show DNA (red line) with immunofluorescence-labeled TRP75 (green lines) (**A, B**), showing TRP75 peaks encasing bacteria DNA peak in the vacuoles in panel **A**, but not in panel **B**. (**C**) Numbers of green puncta on the surface of infected cells (N = 33). ***, *P* < 0.001, unpaired Student *t* test.

### TRP75ΔC and WT *E. japonica* similarly infect cells in culture and mice, and elicit similar disease in mice

A time course of WT and TRP75ΔC *E. japonica* replication in cultured DH82 cells showed similar growth rates based on qPCR (Fig. 4A). Next, we carried out an *in vivo* assessment of infection with *E. japonica* and ehrlichiosis disease progression in immunocompetent mice, by intraperitoneal (*ip*) inoculation with a lethal dose of WT *E. japonica* (approximately 1,000 – 2,000 bacteria) or an equivalent number of TRP75ΔC bacteria with daily monitoring. Both groups of mice developed progressively severe clinical signs (squinty eyes, hunched back, ruffled fur, anorexia, dehydration, inactivity), resulting in rapid severe body weight loss starting at 4-day post inoculation (dpi) and euthanasia at 7 dpi (Fig. 4B). There were no significant differences in *Ehrlichia* load in the blood at 7 dpi between these two groups of infected mice (Fig. 4C). Thus, the C-terminal truncation of TRP75 did not significantly alter *E. japonica* growth in DH82 cells or in mice, or *E. japonica*-caused clinical disease in mice.

**FIG 4.**
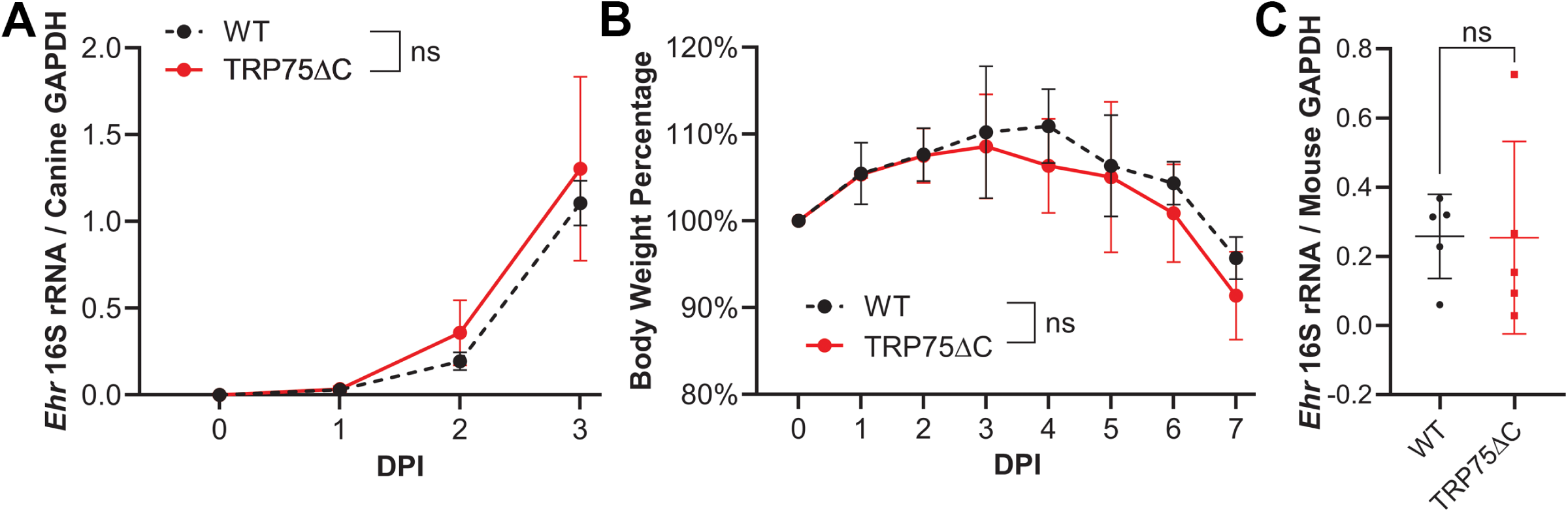
Growth curves of TRP75ΔC and WT *Ehrlichia* and pathogenesis in mice. (**A**) Synchronous growth of TRP75ΔC and WT *E. japonica* in DH82 cells. (**B, C**) Mice were *ip* inoculated with ∼1,000 TRP75ΔC or WT *E. japonica.* (**B**) Body weight was monitored daily, and all mice were moribund at 7 dpi and experiment was terminated/mice were euthanized. (**A, C**) qPCR analysis of cell cultures and infected mouse blood samples using *Ehrlichia* 16S rRNA normalized by canine or mouse GAPDH. Two-way or one-way ANOVA, N = 5; ns, not significantly different.

### Larval and nymphal ticks fed on TRP75ΔC *E. japonica*-infected mice cannot complete molting

Given the similar growth of WT and TRP75ΔC *E. japonica* in cultured ISE6 tick cells (Fig. 1), we next examined if *I. scapularis* ticks were infected similarly with WT and TRP75ΔC *E. japonica* by allowing to feed on infected mice. As shown in Fig. 5, ticks acquired similar amounts of blood from WT and TRP75ΔC *E. japonica-*infected mice as uninfected mice (estimated based on mouse GAPDH DNA levels in ticks) at both larval (Fig. 5A) and nymphal stages (Fig. 5D), and were infected similarly with WT and TRP75ΔC *E. japonica* (estimated based on *Ehrlichia* 16S rRNA gene levels in ticks (larva, Fig. 5B; nymph, Fig. 5E). However, at both larval and nymphal stages, most of the TRP75ΔC *Ehrlichia-*infected larvae and nymphs could not complete molting to the next developmental stage (only ∼10% molted successfully by 10 weeks post-feeding) (larva, Fig. 5C; nymph, Fig. 5F). In contrast, engorged WT *E. japonica*-infected ticks molted to the next stage similarly as engorged uninfected ticks (>80% molted successfully by 10 weeks post-feeding) (Fig. 5C and Fig. 5F). We used 10 weeks post-feeding as cut-off, because we did not observe any additional molting ticks after 10 weeks up to 1.5-year.

**FIG 5.**
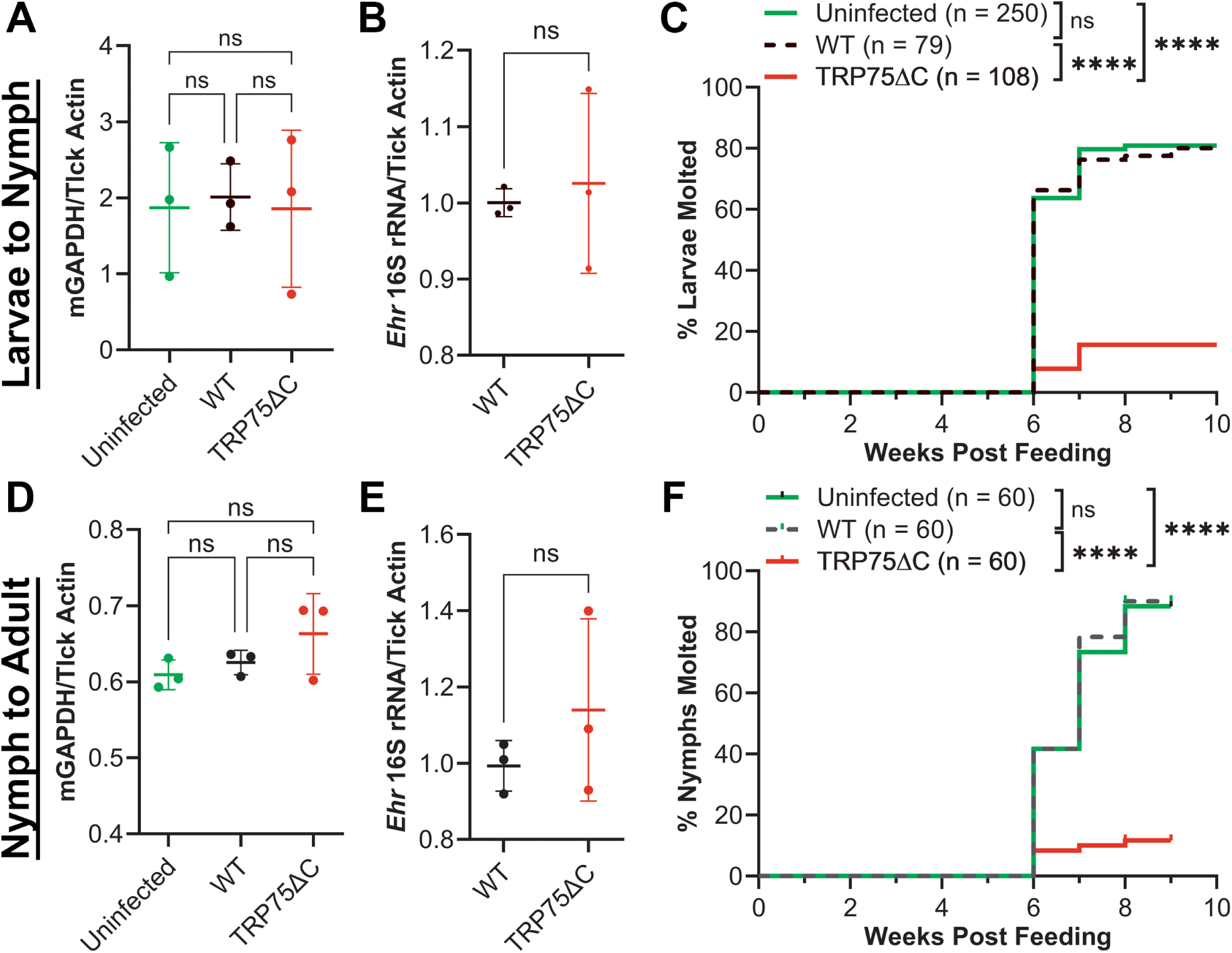
Molting of larval and nymphal ticks fed on WT or TRP75ΔC *E. japonica*-infected, or uninfected mice. *I. scapularis* larvae (**A-C**) and nymphs (**D-F**) were allowed to feed on uninfected mice for 4 d, or WT or TRP75ΔC *E. japonica*-infected mice at 3 dpi and removed at 7 dpi. At 7 dpi, ticks acquired the same amount of blood (*mGAPDH/Tick actin*, **A** and **D**) and *Ehrlichia* (16S rRNA/*Tick Actin*, **B** and **E**). N = 3 of five pooled larvae or three pooled nymphs each. One-way ANOVA; ns, not significant. (**C** and **F**) % molted ticks molted were determined every week post-feeding (week 0). ns, not significantly different; **** significantly different (*P* < 0.0001) by the Mantel-Cox test.

### Off-host post-feeding ecdysone receptor mRNA is upregulated in uninfected and WT *E. japonica*-infected ticks, but is not TRP75ΔC *E. japonica*-infected larval ticks

In order to molt, ticks convert imbibed blood cholesterol into the bio-active molting hormone 20-hydroxyecdysone (20E) by several ecdysone biosynthetic enzymes, including Phantom, Shadow, and Shade (39–41). Ticks synthesize prohormone ecdysone in the integument (epidermis), unlike insects that use a distinct prothoracic gland (40). In both insects and ticks, ecdysone conversion to 20E by Shade occurs in peripheral tissues like the fat body (39, 40). 20E can bind to the nuclear receptor complex, ecdysone receptor (EcR) and Ultraspiracle (Usp), to activate the downstream transcription factors that promote developmental transitions of insects and soft ticks (39, 41). In agreement with the similar molting rates between WT *E. japonica*-infected and uninfected ticks, there was no significant difference in mRNA expression of Phantom, Shadow, and Shade, and EcR from day 0 to 15 post-feeding (Fig. 6). mRNA expression of Phantom was similarly upregulated in uninfected, and WT and TRP75ΔC *E. japonica*-infected ticks from day 0 to 10 post-feeding, and expression plateaued by day 15 (Fig. 6A). In all groups of ticks, Shade expression was relatively unchanged from day 0 to day 10 post-feeding regardless of infection status (Fig. 6B), whereas mRNA expression of Shadow was approximately 100-fold lower in TRP75ΔC *E. japonica*-infected than WT *E. japonica* -infected or uninfected ticks at day 0 post-feeding, and exponential increase occurred to catch up to reach to the similar level as WT-infected and uninfected ticks at day 15 post-feeding (Fig. 6C). Strikingly, mRNA of EcR was significantly upregulated over 1,000-fold in both WT *E. japonica* and uninfected ticks between day 0 and 15 post-feeding, but was not upregulated in TRP75ΔC *E. japonica*-infected ticks (Fig. 6D).

**FIG 6.**
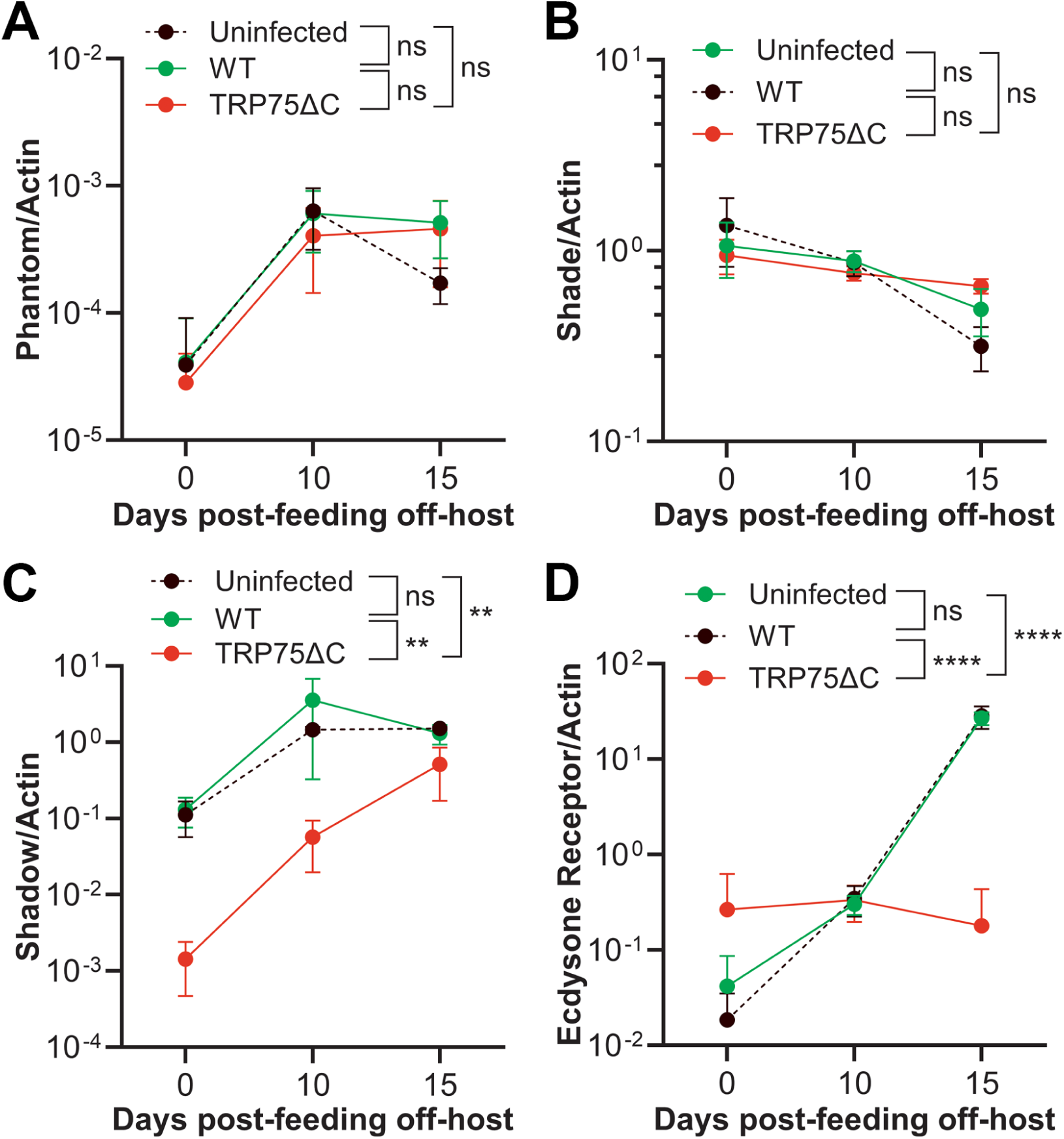
RT-PCR analysis of three 20E synthesis enzymes and EcR expression post-feeding. *I. scapularis* larvae were allowed to feed on uninfected mice for 4 d, or WT or TRP75ΔC *E. japonica-*infected mice at 3 dpi and removed at 7 dpi. (**A-D**) RT-qPCR of three enzymes (phantom, shadow, and shade) in the 20E biosynthesis pathway and EcR/tick actin at 0, 10, and 15 days post-feeding. N=3 of five pooled larvae each. Two-way ANOVA; ns, not significant; ** *P* < 0.01; **** *P* < 0.0001.

### Trans-complementation of TRP75ΔC *E. japonica-*infected nymphal ticks with recombinant TRP75 rescues the molting defect

To determine if the molting defect of TRP75ΔC *E. japonica*-infected ticks was due to a lack of full-length TRP75, we cloned and purified endotoxin-free recombinant full-length TRP75 (rTRP75) (Fig. 7A). Since the TRP75ΔC mutation did not impact the ability of ticks to acquire *E. japonica* (Fig. 5), 160 of engorged nymphs fed on TRP75ΔC *Eja*-infected mice were divided into two groups of 80 nymphs each for microinjection with rTRP75 or phosphate-buffered saline (PBS) as a negative control, within several hours after harvesting ticks. At 1-day post-feeding/microinjection, both groups of ticks were similarly infected and acquired mouse blood (Fig. 7B and C). However, as shown Fig. 7D, the microinjection with rTRP75, but not PBS, significantly increased molting efficiency of nymphs fed on TRP75ΔC *Eja*-infected mice from ∼20% to over 70% at 10 weeks post-feeding.

**FIG 7.**
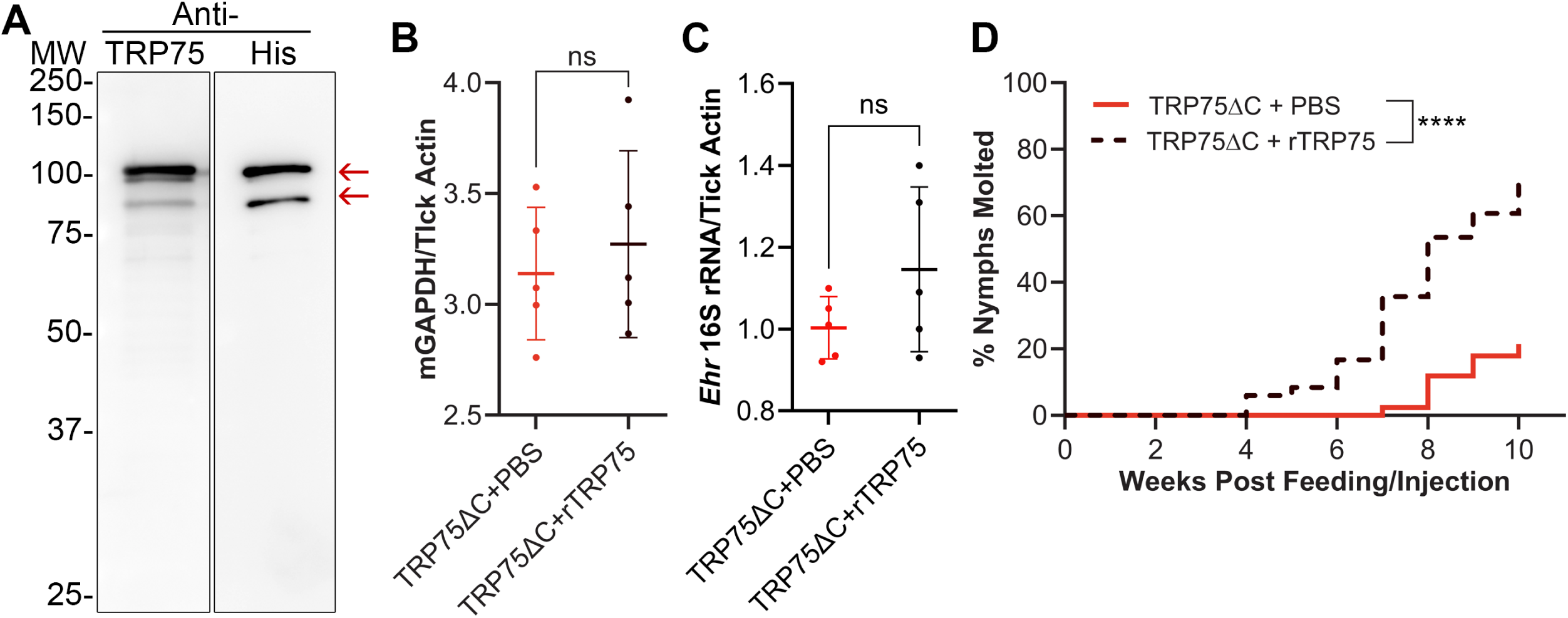
Molting efficiency of TRP75ΔC *E. japonica-*infected nymphs microinjected with rTRP75 or PBS. (**A**) Western blot analysis of affinity-purified recombinant *E. japonica* TRP75 (rTRP75) detected with anti-*E. chaffeensis* TRP75 and anti-His as indicated by red arrows. Lower bands, cleaved TRP75. (**B-D)** Molting efficiency of TRP75ΔC *Ehrlichia*–infected nymphal ticks microinjected with rTRP75 or PBS. *I. scapularis* nymphs were allowed to feed on ten TRP75ΔC *E. japonica*–infected mice at 3 dpi and removed at 7 dpi. Freshly engorged ticks from five infected mice each were microinjected with rTRP75 or PBS (negative control). (**B-C**) At day 1 post-microinjection, imbibed mouse blood levels (**B**) and *Ehrlichia* infection (**C**) were determined by qPCR with mouse *GAPDH* or *Ehrlichia* 16S rRNA normalized with tick *Actin*, respectively. N=5 of three pooled nymphs each. One-way ANOVA; ns, not significant. (**D**) The percentage of nymphs that molted was determined every week post-feeding (post-injection). rTRP75 vs. PBS: significantly different (*P* < 0.0001) by the Mantel-Cox test. N=80 each.

### Microinjection of TRP75ΔC *E. japonica-*infected nymphal ticks with rTRP75 upregulates EcR mRNA

Since microinjection of full-length rTRP75 complements the molting defect of TRP75ΔC *E. japonica*-infected nymphs (Fig. 7), we assessed whether rTRP75 can restore EcR expression in ticks post-feeding. TRP75ΔC *E. japonica*-infected nymphs were microinjected with full-length rTRP75 as described above, and expressions of ecdysone biosynthesis genes were determined by RT-qPCR. From day 0 to 15 post-feeding, there were only minor differences in mRNA expression of Phantom, Shadow, and Shade between rTRP75 and PBS-microinjected TRP75ΔC *Ehrlichia*-infected nymphs (Fig. 8). Between days 1 and 10 post-feeding, these three ecdysone synthesis genes were upregulated, and leveled off at day 15 post-feeding. However, EcR mRNA was significantly upregulated in nymphs microinjected with rTRP75 compared with PBS-microinjected nymphs at day 10-15 post-feeding (Fig. 8).

**FIG 8.**
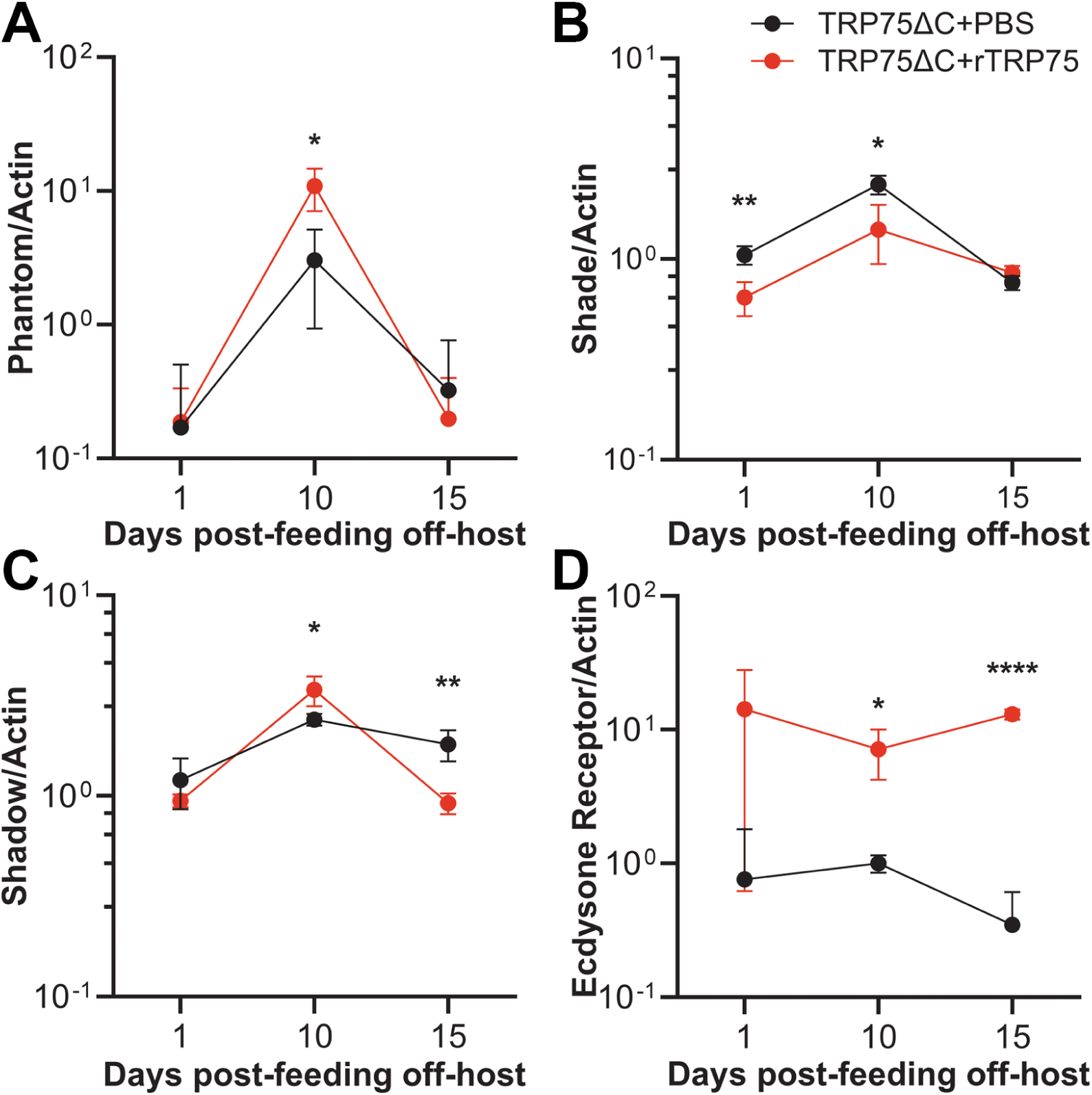
20E synthesis enzymes and EcR transcription in TRP75ΔC *Ehrlichia*-infected ticks upon rTRP75 microinjection. Each of 15 TRP75ΔC *E. japonica*–infected *I. scapularis* nymphs microinjected with rTRP75 or PBS. (**A–D**) RT-qPCR of three enzymes of the 20E biosynthesis pathway (phantom, shadow, and shade) and EcR/tick actin at days 1, 10, and 15 post-feeding off-host. N = 5 of three pooled nymphs each. 2-way ANOVA where ns, not significant, * *P* < 0.05, ** *P* < 0.01, **** *P* < 0.0001.

## DISCUSSION

Our results demonstrate that the C-terminus of *Ehrlichia* TRP75 has a critical role in survival of *Ehrlichia* and its tick hosts by allowing infected ticks to molt into the next developmental stage as effectively as uninfected ticks. Molting of infected ticks is essential for transstadial transmission of *Ehrlichia,* thereby allowing mammalian transmission during the next blood meal. The successful trans-complementation of the molting defect of TRP75ΔC *Ehrlichia*-infected ticks upon rTRP75 microinjection suggests that TRP75 is sufficient to rescue the tick molting defect. To our knowledge, this is the first demonstration of complementation in ticks by microinjection of the any recombinant bacterial protein. Thus, the full-length TRP75 in *Ehrlichia* is required for successful molting of infected ticks, although TRP75 is not required for molting of uninfected ticks, suggesting that TRP75 in *Ehrlichia* may neutralize or compensate for the infection-induced deficit or stress in ticks.

In post-feeding engorged ticks, ecdysone is synthesized in the integument and EcR is synthesized in peripheral tissues (39, 40). Our results show that, compared with uninfected ticks and each of WT and TRP75ΔC *Ehrlichia*–infected ticks, the temporal transcription pattern of the three ecdysone synthesis genes we examined during the post-feeding/pre-molting period did not differ substantially. However, EcR up-regulation in WT was suppressed in TRP75ΔC *Ehrlichia*-infected ticks. These results indicate that full-length TRP75 or TRP75 IDR is required for EcR upregulation in *Ehrlichia*-infected larval and nymphal ticks.

Although *E. chaffeensis* TRP75 has been suggested to be an effector of the Type I secretion system, unlike *E. chaffeensis* TRP120, this has not been demonstrated in a heterologous *E. coli* Type 1 secretion system (42). However, the *E. chaffeensis* TRP75 fragment (aa 23 to 583), lacking lipobox was reported to interact with 90 different human proteins based on results of a yeast two-hybrid analysis and 13 selected human proteins were shown to colocalize with TRP75 based on antibody staining and co-immunoprecipitation (34). Additionally, in *E. chaffeensis*-infected THP-1 cells, siRNA knockdown of several of these human proteins significantly reduced *E. chaffeensis* infection of THP-1 cells, suggesting that TRP75 regulates human cell apoptosis, cytoskeleton reorganization, and/or evasion of host immune defense (34). However, whether TRP75 knockdown or knockout in *Ehrlichia* inhibits *Ehrlichia* infection is unknown. A recent study reported that transfection of HEK293 cells with GFP-TRP75 facilitates host STAT3 phosphorylation and nuclear translocation (35). In contrast, nothing is known about the roles of TRP75 in *Ehrlichia* infection in tick cells and ticks, or in mammalian infection, or tick transmission *in vivo*. Thus, the present study fills a void in knowledge concerning the roles of *Ehrlichia* TRP75,—specifically its C-terminal IDR—*in vitro* and *in vivo*, suggesting its essential roles in ticks *in vivo*.

The EcR is a nuclear hormone receptor that forms a functional heterodimer with the retinoid X receptor (RXR/Usp) to mediate 20E signaling that is critical for tick development, molting, and reproduction (41). Tick EcR acts downstream or in conjunction with systemic pathways, such as those governed by the tick lipid-binding protein croquemort, which impacts ecdysteroid synthesis required for successful molting and feeding (43). However, rTRP75 rescued the molting defect in TRP75ΔC *Ehrlichia*-infected ticks with minimal impact on the mRNA levels of 3 enzymes involved in 20E synthesis during the pre-molting/post-feeding period, in contrast to its dramatic impact on EcR, suggesting that the mechanism of TRP75 may differ from that of croquemort with respect to rescue molting defect.

We previously showed that knockdown of TRP120 in *E. japonica* neither inhibits *Ehrlichia* infection in tick or mammalian host cells in culture or inhibits mouse infection or lethal ehrlichiosis in mice (30). However, knockout of TRP120 almost completely abrogates bacteremia (ehrlichemia) without inhibiting tissue infection in mice, thereby blocking tick acquisition of *Ehrlichia* upon blood feeding on the infected mice (30). Thus, TRP120 is essential for *Ehrlichia* survival by facilitating tick-mediated transmission of *Ehrlichia*, a core aspect of the natural *Ehrlichia* life cycle (30). TRP75 is the second molecule found to be required for tick-to-mammal transmission, suggesting that both TRP molecules in tick-borne *Ehrlichia* species have evolved for *Ehrlichia* survival at the vulnerable transition stage between ticks and mammals and vice versa. Because these two TRP molecules or their fragments are essential for the natural life-cycle of *Ehrlichia-*infected ticks, they could potentially be exploited to selectively eliminate *Ehrlichia-*infected ticks, and thereby prevent *Ehrlichia* infection and ehrlichiosis in humans and animals.

## MATERIALS AND METHODS

### Ethics statement

All animal experiments were performed in accordance with guidelines of the Ohio State University Institutional Animal Care and Use Committee, which also approved the experimental protocol. The university program has full accreditation by the Association for Assessment and Accreditation of Laboratory Animal Care International under 000028 and has Public Health Services assurance renewal A3261-01. The program is licensed by the USDA, number 31-R-014, and is in full compliance with Animal Welfare Regulations.

### Culture of host cells and their infection with *Ehrlichia*

Uninfected DH82 cells were cultured in DMEM (Dulbecco minimal essential medium; Mediatech, Manassas, VA) supplemented with 5% fetal bovine serum (FBS; Atlanta Biologicals, Lawrenceville, GA) and 2 mM ʟ-glutamine (ʟ-Gln; Gibco, Waltham, MA). DH82 cells infected with WT *E. japonica* strain HF (17) or TRP75ΔC mutant (H130-A1) were cultured with the addition of 0.1 μg/ml cycloheximide (Sigma, Burlington, MA). Uninfected and infected RF/6A cells (ATCC, Manassas, VA) were cultured in DMEM supplemented with 5% FBS and 2 mM ʟ-glutamine. Both DH82 and RF/6A cells were cultured at 37°C under 5% CO_2_ in a humidified atmosphere. Uninfected and infected ISE6 cells were cultured in L15C300 medium as described (44, 45). Bacterial infection levels were determined by HEMA3 staining (Thermo Fisher Scientific, Waltham, MA) and microscopical evaluation. Synchronous culture of WT and TRP75Δ *E. japonica* was done by inoculating DH82 cells at multiplicity of infection (MOI) of 100-200 and harvesting at 1-3 dpi as previously described (30, 46), and infectivity were quantified by qPCR.

### Clonal isolation of TRP75ΔC mutant and confirmation of single Himar1 insertion

The *E. japonica* Himar1 mutant pool H130-A1 was originally generated and genetically mapped as described previously (31). For clonal isolation, DH82 cells were seeded in flat-bottom 96-well plates (Greiner, Monroe, NC) at 10⁴ cells per well in 50 μl of DMEM and inoculated with approximately two H130-A1–infected DH82 cells per well. After overnight incubation, 150 μl of fresh culture medium was added to each well, and the cultures were grown for 6 days to reach full confluence. Cells were detached using sterile pipette tips, and 20-μl aliquots of each cell suspension were transferred to corresponding wells in two replicate 96-well plates. Each well received 180 μl of fresh culture medium, and the cultures were grown for an additional 6 days to reach full confluence and allow further proliferation of the mutant to increase infectivity. To identify wells containing the mCherry-positive mutant, the medium was removed from one replicate plate, and 50 μl of 1× PBS was added to each well. Cells were detached using sterile pipette tips and thoroughly resuspended. A 1-μL aliquot of each suspension was used directly as the template for PCR with mCherry-specific primers and Platinum Direct PCR Universal Master Mix (Invitrogen, Carlsbad, CA), following the manufacturer’s instructions. Wells yielding an amplification product detected by agarose gel electrophoresis were designated mCherry-positive. Cultures from the corresponding wells of the second replicate plate were expanded and assessed for mutant clonality by PCR using primers flanking the insertion site at genomic position 556,130 (Fig. 1). Himar1 insertion copy number per genome was estimated by qPCR targeting mCherry and the *Ehrlichia* 16S rRNA gene in DNA isolated from TRP75ΔC mutant clones. The plasmids pCis-mCherry-SS Himar A7 and pUC19 containing the *Ehrlichia* 16S rRNA gene were used as standards for absolute quantification of mCherry and 16S rRNA gene copy numbers, respectively (31). Insertion copy number per genome was calculated as the ratio of mCherry to 16S rRNA gene copy numbers. All primers used in this study are listed in Table S1.

### Amino-acid sequence analysis and structure prediction

The amino-acid sequence of *E. japonica* TRP75 was analyzed by the program Protean (DNASTAR Lasergene) (47), and the three-dimensional structure diagram was rendered by the AlphaFold algorithm (https://alphafold.ebi.ac.uk/entry/X5H2X1) (48). SignalP 6.0 and DeepLocPro 1.0 (https://services.healthtech.dtu.dk/) (49) was used to predict protein localization and lipobox domains.

### Western blotting and immunofluorescence labeling

TRP75ΔC or WT–*Eja*-infected DH82 cells were examined for TRP75 protein expression by western blotting using a dog antiserum against recombinant *E. chaffeensis* TRP75 (33), rabbit anti-*E. chaffeensis* P28 (38), and rabbit anti-β-actin (Sigma-Aldrich, St. Louis, MO). Infected and uninfected DH82 cells were lysed in MPER (Pierce Biotech, Rockford, IL) w/ protease inhibitor cocktail (Calbiochem, San Diego, CA) (5 × 10^6^ cells in 500 µl), mixed with 2× SDS-PAGE sample buffer, and boiled for 5 min. Following SDS-PAGE, samples were subjected to semi-dry transfer to a nitrocellulose membrane for 1 h. Dog anti-*E. chaffeensis* TRP75 serum (1/400 dilution) was pre-adsorbed in a nitrocellulose membrane with 10 lanes of uninfected DH82 cell lysates for 2 h at room temperature, then incubated with *Ehrlichia* sample membrane overnight at 4°C.

Cloned TRP75ΔC or WT *E. japonica*–infected RF/6A cells were immunofluorescence labeled with dog anti-*E. chaffeensis* TRP75 and FITC-conjugated rabbit anti-dog IgG (Jackson ImmunoResearch, West Grove, PA). The cell-permeant Hoechst 33342 (Thermo Fisher Scientific) was used to stain nucleic acids. Fluorescence images with overlay differential interference contrast images (DIC) were acquired and analyzed with a Leica THUNDER imaging system with computational clearing to remove any out-of-focus background (Leica Microsystems, Deerfield, IL). NIH ImageJ software was used to measure line scan relative fluorescence intensities.

### Cloning of rTRP75

The gene encoding *E. japonica TRP75* (EHF_0506) (22 – 684 aa) was PCR amplified and cloned into pET-33b(+) (Novagen, Gibbstown, NJ) at the EcoRI and XhoI restriction sites to create a plasmid expressing N-terminal 6×His-tagged TRP75 (rTRP75). rTRP75 protein was affinity purified from soluble fractions of transformed *E. coli* BL21-CodonPlus (DE3)-RIPL competent cells (Agilent Technologies, Santa Clara, CA) using HisPur Cobalt Resin (Thermo Fisher) as described (50). Potential endotoxin was removed with Pierce High-Capacity Endotoxin Removal Resin (Thermo scientific, Rockford, IL), and verified by the ToxinSensor Chromogenic LAL endotoxin assay kit (GenScript, Piscataway, NJ) to be < 0.01 EU/ml.

### Inoculation of mice and sample collection

ICR mice at age 6 weeks old (Envigo, Indianapolis, IN) (two groups of 5 mice each) were *ip* inoculated with infected DH82 cells containing approximately 1,000 – 2,000 WT or TRP75ΔC *E. japonica* resuspended in 0.5 ml serum-free DMEM using a 1-ml Tuberculin syringe and 26-gauge needle (BD, Franklin Lakes, NJ). Clinical signs and body weight of individual mouse were monitored daily. Blood samples were collected in Covidien Monoject with 0.04 ml of 7.5% EDTA Liquid Additive tubes (Thermo Fisher Scientific) by cardiac puncture, and stored at –20°C prior to DNA extraction by using DNeasy Blood & Tissue kits (Qiagen, Germantown, MD).

### Tick attachment and off-host molting assay

Three 8-week-old C57BL/6 mice (originally from Inotiv; West Lafayette, IN; and bred at OSU animal facility), each were *ip* inoculated with 2,000 – 3,000 TRP75ΔC or WT *E. japonica*-infected DH82 cells. Three uninfected mice were used as the control. At 3 d pi, *I. scapularis* larval tick mass or approximately 25 nymphal ticks (Oklahoma State University Tick Rearing Facility; Stillwater, OK) were placed inside of a tick confinement chamber glued onto the shaved back of each mouse under inhalational anesthesia as described (30). At day 4 post tick-attachment (7 dpi), 50-150 engorged larval or ∼25 nymphal ticks were recovered from each mouse, and incubated off-host at 22°C and 99% relative humidity with a 10 h dark/14 h light cycle in the environmental chamber. At day 0 post-feeding five engorged larval and three engorged nymphal ticks from each mouse were pooled and frozen at −20°C until DNA extraction. At day 0, 10, and 15 post-feeding, five engorged larval and three engorged nymphal ticks from each mouse were pooled, bisected and preserved in RNAlater buffer (Thermo Fisher Scientific) overnight and transferred to –80°C until RNA extraction. Remaining ticks were kept in the incubator for 10 weeks to 1.5 year, and % molted/total ticks and molted week range were scored.

### Complementation of TRP75ΔC mutant-infected ticks with rTRP75

At 3 dpi with TRP75ΔC *E. japonica*-infected DH82 cells, 25-35 nymphs were fed on each of 10 mice and removed at day 4 post tick-attachment (7 dpi). Engorged ticks from 5 mice each were microinjected with 1 µl of 10-15 ng/µl of recombinant TRP75 (rTRP75) in PBS, or PBS alone, using Hamilton Model 62 RN Syringe with a 33-G needle (Hamilton, Reno, NV) under dissection microscope (Autofocus 4K Digital Microscope. Model: EM4K-AF 8-inch (Elikliv, Guandong, China). Then ticks were incubated off-host at 22°C and 99% relative humidity with a 10 h dark/14 h light cycle in the environmental chamber. At day 1 post-feeding three engorged nymphal ticks from each mouse were pooled and frozen at −20°C until DNA extraction. At day 1, 10, and 15 post-feeding, three engorged nymphal ticks from each mouse were pooled, bisected and were preserved in RNAlater buffer (Thermo Fisher Scientific) overnight and transferred to –80°C until RNA extraction. Remaining ticks were kept in the incubator for 10 weeks, and % molted/total ticks and molted week range were compared between rTRP75-injected and PBS-infected nymphs.

### qPCR and RT-qPCR

Tick samples from each mouse stored as above were homogenized by using tissue grinders (Sigma-Aldrich) and processed for DNA extraction by using DNeasy Blood & Tissue kits (Qiagen). For RNA extraction, tick samples were similarly homogenized, and RNA was isolated using RNeasy Mini kits (Qiagen). cDNA was synthesized from 1–2 µg of extracted RNA using the Maxima H minus First Strand cDNA synthesis kit (Thermo Fisher Scientific) with random hexamer primers. qPCR and RT-qPCR analysis were performed using Maxima SYBR Green/ROX Master Mix (Thermo Fisher Scientific) according to the manufacturers’ protocols in Mx3000P (Stratagene, La Jolla, CA) or AriaMx Real-time PCR system (Agilent, Santa Clara, CA). To quantify the exact inoculums of *E. japonica*, an absolute quantification method was used by creating a standard curve of *Ehrlichia* 16S rDNA cloned into plasmid pUC19 as a standard. To quantify relative *E. japonica* loads in mouse blood, or ticks, qPCR was performed using primers specific for *Ehrlichia* 16S rDNA and mouse *GAPDH* or *I. scapularis Actin* gene (51). To assess tick gene expression, RT-qPCR was performed using primers specific for *I. scapularis* Shadow, Shade, Phantom, and EcR, along with *I. scapularis* Actin for normalization using primers listed in Table S1.

### Statistical analysis

The significance of differences between values was assessed using the two-tailed Student *t* test or analysis of variance (ANOVA). Data analysis and figure drawing were performed with Prism 9 software (GraphPad, San Diego, CA). The Mantel-Cox test, a nonparametric, chi-square-based method was used to compare the molting distributions of 3 or 2 independent groups over time by Prism 11 software (GraphPad). *P* value of <0.05 was considered to reflect statistical significance.

## Acknowledgements

Jeffery Kawahara is the recipient of an OSU CVM 2024 C. Glenn Barber fellowship. This work was partially supported by the National Institutes of Health grant R01 AI152223 to YR from the National Institutes of Health.

