## Supplemental Information for "Recombinant *Ehrlichia* TRP75 Rescues Molting Failure of Ticks Infected with TRP75 Mutant *Ehrlichia*"

**Running Title**: *Ehrlichia* TRP75 prevents molting failure of infected ticks

*Equally contributed.

### Corresponding author: Department of Veterinary Biosciences, College of Veterinary Medicine, The Ohio State University, 1925 Coffey Road, Columbus, OH 43210-1093. Phone: (614) 292-9677. Fax: (614) 292-6473.

The authors declare no competing interests.

#### Supplemental Materials

Supplemental Materials and Methods, and Table S1 can be accessed on the publisher's website.

**Supplemental Table S1. Primers used in this study.**

| **Target** | **Primer sequence (5'-3')** | **Notes** |
|---|---|---|
| **TRP75 cloning †** | | |
| *E. japonica TRP75* | F: atc GAATTC tTCAAATAAGAGTAGATATCAGTTTAGC<br>R: agt CTCGAG **TTA**ATCATTATAGTAATAAGTAATACTG | For cloning *E. japonica TRP75* into *EcoRI* (F) and *XhoI* (R) sites in pET33b (+) plasmid* |
| **Flanking PCR** | | |
| *E. japonica* i-556130 | F: AAGACAAAGGGTTGCGTTCA<br>R: ACTTGACCATTGTGCTCAGATAC | For flanking the Himar1 insertion site i-556130 of *E. japonica* TRP75ΔC (31) |
| **RT-qPCR or qPCR** | | |
| *Ehr* 16S rRNA | F: CGGGGGAAAGATTTATCGCTATTA<br>R: CGCTTGCCCCCTCCGTATTA | For qPCR and RT-qPCR to quantify *Ehrlichia* (52) |
| Mouse GAPDH | F: GTTGTCTCCTGCGACTTCA<br>R: GGTGGTCCAGGGTTTCTTA | For qPCR and RT-qPCR to normalize quantification in mouse (53) |
| *I. scapularis* actin | F: GCCCTGGACTCCGAGCAG<br>R: CCGTCGGGAAGCTCGTAGG | For qPCR to normalize quantification in *I. scapularis* (54) |
| *I. scapularis* actin | F: CTTGGAGGGAGACATCTTTGTG<br>R: CGCTCCATGTCATTCCAATCT | For RT-qPCR to normalize quantification in ISE6 cells (51, 55) |
| Himar1 mCherry | F: CTCCGGGTACATTCTCTCGC<br>R: GGAACGCGTCATGAACTTCG | For qPCR to quantify the copy number of Himar1 insertion in *E. japonica* (30) |
| ***I. scapularis* Ecdysone pathway** | | |
| Phantom | F: GACTCTAGCCGGAGTGTACG<br>R: GAAGACGCGATGCAGGAAAG | For qPCR and RT-qPCR * |
| Shade | F: TACTTCCGATGGGGACGCTAC<br>R: GTGTGGGTGTTCGGTTCGAT | For qPCR and RT-qPCR (43) |

| | | |
|---|---|---|
| Shadow | F: TCCGGGAAGCTCTACGGTTA<br>R: ACGAGTTGCTGCTCTGACTC | For qPCR and RT-qPCR * |
| Ecdysone receptor | F: ATGAAGCCCGTGCTGCTG<br>R: GACGGCGACGAGATCCAG | For RT-qPCR * |

* Primers were designed in this study.
† F, forward; R, reverse complement primers; underlined sequences, restriction enzyme sites; bold, stop codon.

#### Supplemental figures

**FIG. S1. Amino-acid sequence alignment of *E. japonica* HF TRP75C with *E. chaffeensis* TRP75C.**

The C-terminal amino-acid sequence of *E. japonica* TRP75 (*Eja*TRP75C, 113 aa downstream of the Himar1 insertion: 572–684 aa; full length: 684 aa) was aligned with *Ech* TRP75C (C-terminal 111 aa: 465–575 aa; full length: 583 aa) using Blastp, revealing 72% sequence similarity.

#### Figure S1

```
Score = 124 bits (312), Expect = 3e-31
Identities = 64/116 (55%), Positives = 84/116 (72%), Gaps = 8/116 (6%)

EjaTRP75C: 572   NDEEPVLSQSRHDLEPAKLISEGSEQKNNYKAVHFLSFLKEQNKDQNDTTEELQKAENKN   631
EchTRP75C: 465   DSEEPTFSQPKSDVESPKQVSESDEQKNSHKAVHFLSFLNEPSEDQNDTTEELQNTEDKK   524

EjaTRP75C: 632   D---QENAEVSEHNGQVNIGNDGIMSEEEKDYVLLQRIEGMKEYDEDYSITYYYND   684
EchTRP75C: 525   DNLLEENVKISEKDDQ-----QVVISIEEENQMLLQSIKKMKEYDEDYSITYYYDD   575
```
